# An accurate and rapid continuous wavelet dynamic time warping algorithm for unbalanced global mapping in nanopore sequencing

**DOI:** 10.1101/238857

**Authors:** Renmin Han, Yu Li, Sheng Wang, Xin Gao

## Abstract

Long-reads, point-of-care, and PCR-free are the promises brought by nanopore sequencing. Among various steps in nanopore data analysis, the global mapping between the raw electrical current signal sequence and the expected signal sequence from the pore model serves as the key building block to base calling, reads mapping, variant identification, and methylation detection. However, the ultra-long reads of nanopore sequencing and an order of magnitude difference in the sampling speeds of the two sequences make the classical dynamic time warping (DTW) and its variants infeasible to solve the problem. Here, we propose a novel multi-level DTW algorithm, cwDTW, based on continuous wavelet transforms with different scales of the two signal sequences. Our algorithm starts from low-resolution wavelet transforms of the two sequences, such that the transformed sequences are short and have similar sampling rates. Then the peaks and nadirs of the transformed sequences are extracted to form feature sequences with similar lengths, which can be easily mapped by the original DTW. Our algorithm then recursively projects the warping path from a lower-resolution level to a higher-resolution one by building a context-dependent boundary and enabling a constrained search for the warping path in the latter. Comprehensive experiments on two real nanopore datasets on human and on *Pandoraea pnomenusa*, as well as two benchmark datasets from previous studies, demonstrate the efficiency and effectiveness of the proposed algorithm. In particular, cwDTW can almost always generate warping paths that are very close to the original DTW, which are remarkably more accurate than the state-of-the-art methods including Fast-DTW and PrunedDTW. Meanwhile, on the real nanopore datasets, cwDTW is about 440 times faster than FastDTW and 3000 times faster than the original DTW. Our program is available at https://github.com/realbigws/cwDTW.

## 1 Introduction

DNA sequencing has been dominated by sequencing-by-synthesis technologies for decades [1]. Nowadays, singlemolecule sequencing based on nanopore technologies has emerged with the promises of long-reads, point-of-care, and PCR-free [2]. Long-reads provides great potentials for *de novo* transcriptome analysis, which is able to span more repetitive regions and multiple exon junctions [3]; point-of-care makes it possible for the sequencing to be conducted immediately at anywhere in real-time [4]; and PCR-free allows the direct identification of epigenetics [5].

The key innovation of nanopore sequencing is the direct measurement of the changes in the electrical current signal (denoted as the *raw signal*) when a single-strand DNA passes through the nanopore [6] (Fig. 1). Without the needs for polymerase chain reaction (PCR) amplification, nanopore sequencing generates extremely long reads, typically ranging from 12k to 120k bp. At each time point, there are *k* consecutive nucleotides in a pore (denoted as a *k*-mer, where *k* is often 5 or 6). The electrical current signal is measured for each time point of the pore. A *pore model* describes the expected electrical current values for different *k*-mers. In nanopore sequencing, the frequency of the electrical current measurements and the speed of the DNA sequence passing through the pore are not coordinated, which causes the main technical difficulty for nanopore data analysis. In practice, the frequency of the electrical current measurements is 7-9 times higher than the passing speed of the DNA sequence, resulting in an order of magnitude difference in the sampling rates of the raw signal sequence and the expected signal sequence from the pore model.

**Figure 1:**
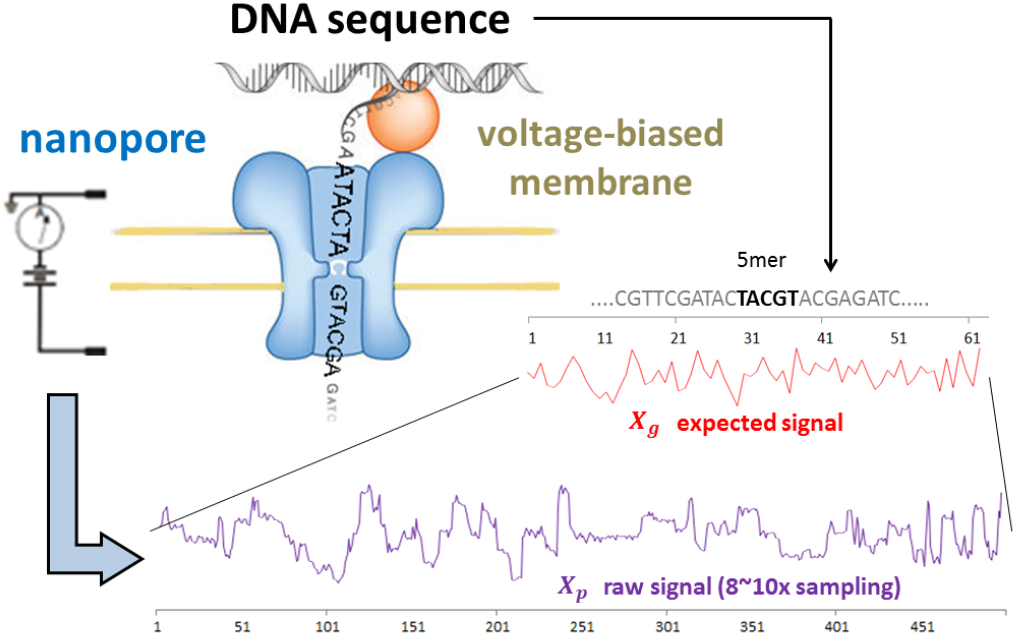
Upper part: pore chemistry contains the voltage-biased membrane embedded with nanopores, in which voltage can be applied to drive the DNA sequence through the pore and electrical current signals (i.e., raw signals) can be measured. Lower part: unbalanced global mapping between the raw signal sequence and the expected signal sequence derived from the DNA sequence and the pore model.

Among various steps in nanopore data analysis, the global mapping between the raw electrical current signal sequence and the expected signal sequence from the pore model serves as the key building block to base calling [7], reads mapping [8], variant identification [9], and methylation detection [5] (the lower part of Fig. 1). Dynamic time warping (DTW) is the most widely-used technique that finds an optimal mapping between two temporal sequences that vary in speed. In DTW, the sequences are warped non-linearly by stretching or shrinking along the time axis [10]. The original DTW has an *O*(*L*_1_*L*_2_) time- and memory-complexity, where *L*_1_ and *L*_2_ are the lengths of the two sequences to be mapped. This complexity severely limits DTW’s applications in various problems that have long sequences, such as nanopore sequencing. To accelerate the analysis of long sequences, a variety of improved DTW have been proposed, which can be roughly categorized into three classes: constrained DTW, coarsening DTW, and multi-level DTW. Constrained DTW (e.g., SparseDTW [11] and PrunedDTW [12]), whose accuracy lies on the strategy of bounding, casts an arbitrary or a predefined boundary to reduce the search space. Coarsening DTW (e.g., PDTW [13], IDDTW [14] and COW [15]) speeds up DTW by operating on a reduced representation of the signals, which is often produced by down-sampling or equal averaging, and then projecting the low-resolution warping path to the full resolution. Yet, the calculated final warping path becomes increasingly inaccurate as the level of coarsening increases. Multi-level DTW (e.g., FastDTW [10] and MultiscaleDTW [16, 17]) combines the ideas of constrained and coarsening DTW. It recursively projects a solution from a low-resolution representation generated by coarsening DTW and refines the projected warping path in high-solution via constrained DTW. Nevertheless, all aforementioned variants of DTW have high risks of failure when the input sequences are noisy and have unbalanced sampling rates, and none of the existing DTW variants can achieve a good balance between accuracy and efficiency on mapping extremely long sequences with unbalanced scales.

In this paper, we propose a novel dynamic time warping algorithm, cwDTW, based on continuous wavelet transform (CWT), to cope with the unbalanced global mapping between two ultra-long signal sequences. The key idea of cwDTW is to obtain a series of highly representative coarsening signals at different resolution levels via CWT. Thus, at each resolution level, the transformed coarsening signal sequences from the two input sequences with unbalanced lengths would have comparable lengths and similar shapes. The warping path obtained from a coarser resolution is used to obtain a stable and narrow context-dependent boundary to constrain the warping path at a refiner resolution. Through this iterative process, our algorithm achieves the global mapping of the input signal sequences in a coarse-to-fine manner.

To our knowledge, it is the first time that continuous wavelet is introduced to the nanopore sequencing analysis and combined with dynamic time warping. Our algorithm benefits from the continuous scale analysis from CWT and is able to utilize the highly representative information embedded in the input signals. Our algorithm has an approximate *O*(*N*) time- and space-complexity, where *N* is the length of the longer sequence, and substantially advances previous methods in terms of the mapping accuracy. Comprehensive experimental results demonstrate the efficiency and effectiveness of cwDTW. Furthermore, cwDTW is not only able to align the raw and expected signal sequences in nanopore, but also applicable to other temporal sequence mapping problems based on biological, video, audio and graphical data.

## 2 Related works

In the section, we provide a brief introduction to continuous wavelet transform and dynamic time warping, which are closely related to the proposed algorithm.

### 2.1 Continuous Wavelet Transform

In mathematics, a continuous wavelet transform (CWT) is used to divide a continuous-time function into wavelets. In particular, the CWT of a one-dimensional signal *X*(*t*) at a scale *a* ∈ ℝ^+^ and translational value *b* ∈ ℝ, denoted as *X_a,b_*, is expressed by the following integral:

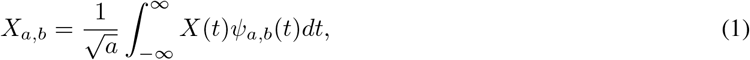

where 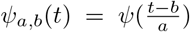 is the mother wavelet which is a continuous function in both the time domain and the frequency domain. In our algorithm, the Mexican hat wavelet, *ψ*(*t*) = (1 − *t*^2^) exp^−*t*^2^/2^, is the default option, but other wavelet functions are also applicable [18].

### 2.2 Dynamic Time Warping

The dynamic time warping (DTW) for mapping two input signal sequences is stated as follows: Given two input signal sequences *X* = *x*_1_, *x*_2_, …, *x*_*L*_1__ and *Y* = *y*_1_, *y*_2_, …, *y*_*L*_2__ of length *L*_1_ and *L*_2_ respectively, construct a warping path *W* = *w*_1_, *w*_2_, …, *w_L_* to minimize the distance measurement 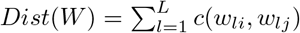, where *L* is the length of the warping path and *c*(*w_li_, w_lj_*) is the Euclidean distance of the *l*th aligned element between the two signal points *x_i_* and *y_j_*. To determine the optimal path *W*, an (*L*_1_ × *L*_2_) matrix *D* is recursively computed, in which the matrix entry *D*(*n, m*) is the total cost of an optimal path between *X* and *Y*. Here *D*(*i, j*) = *min*{*D*(*i* − 1, *j* − 1), *D*(*i, j* − 1), *D*(*i* − 1, *j*)} + *c*(*i, j*) and *c* is the distance between elements *x_i_* in *X* and *y_j_* in *Y*. *D*(*n, m*) can be exactly solved by dynamic programming, resulting in the globally optimal mapping.

Though DTW has been well-established, the original DTW has *O*(*L*_1_*L*_2_) time complexity and needs a matrix *D* with *L*_1_ × *L*_2_ dimension, which is too inefficient and memory-costly for long sequences, such as the ones from nanopore sequencing. To apply DTW in challenging applications, various versions of improved DTW have been proposed, such as FastDTW [10], PrunedDTW [12], SparseDTW [11], and MultiscaleDTW [16, 17].

## 3 Methods

Fig. 2 shows the main workflow of the proposed continuous wavelet dynamic time warping (cwDTW). Three key components are involved: CWT representation, context-dependent constrained DTW, and multi-level refinement. (i) CWT representation is the initial step that runs a continuous wavelet transform on each input signal sequence to obtain an informative representation, followed by peak and nadir picking to produce the low-resolution signals with reduced lengths. (ii) Context-dependent constrained DTW takes a warping path calculated at a lower resolution and determines the search boundary of the path at a higher resolution. (iii) Multi-level refinement combines the low-resolution and high-resolution information from CWT with different scales, and gradually refines the warping path when the level becomes finer and finer, until the final path at the original resolution of the input sequences.

**Figure 2:**
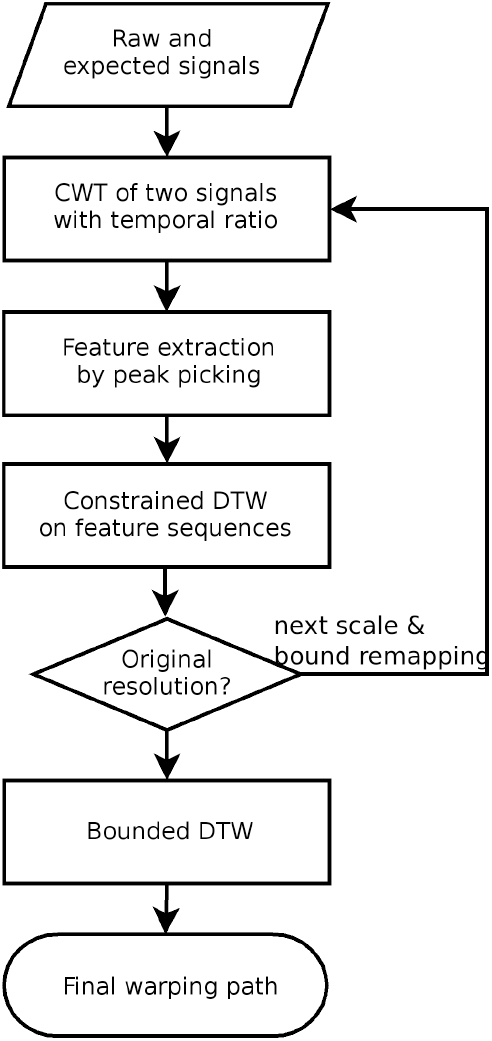
Workflow of the proposed dynamic time warping algorithm based on continuous wavelet transform (cwDTW).

### 3.1 Feature Representation of CWT Spectra

When handling long signal sequences, down sampling combined with multi-scale analysis is widely used to decrease the complexity [10, 16, 17]. Compared with down sampling, wavelet representation is naturally more proper for multiscale analysis [19], where both discrete wavelet transform and continuous wavelet transform are options. Although there have been studies that combine discrete wavelet transform with dynamic time warping [20, 21, 22], such methods have clear limitations.

Discrete wavelet transform is an orthogonal wavelet analysis, in which the number of convolutions at each scale is proportional to the width of the wavelet basis at that scale. To apply discrete wavelet, an alignment to the power of 2 is necessary [21], which usually requires the padding of the signal. This produces a wavelet spectrum that contains discrete “blocks” of wavelet power and is useful for signal processing as it gives the most compact representation of the signal. However, for temporal data analysis, an aperiodic shift in the time series produces a different wavelet spectrum. On the contrary, nonorthogonal transform such as continuous wavelet transform (CWT) is highly redundant at large scales, where the wavelet spectra at adjacent scales are highly correlated. CWT is thus more useful for time series analysis, where smooth and continuous wavelet transforms are expected.

#### 3.1.1 Multi-level representation of CWT

Coming back to Eq.(1), intuitively, the transformed wavelet spectrum *X_a,b_* reflects the pattern matching between the input signal *X* and the wavelet function *ψ*. By changing the scale parameter *a*, larger values correspond to lower frequency signals whereas smaller values correspond to higher frequency signals.

Fig. 3(A) shows the CWT spectra of an expected signal sequence^1^ *X* with different values of the scale parameter *a*. For convenience of analysis, we fix the translational value b as the same index correspondence as *X*. That is, the transformed signals (spectrum) have the same length and retain peer-to-peer index to *X*. Here we use CWT(*X, a*) to denote the transformed spectrum of *X* with the scale parameter *a*. In Fig. 3(A), the input is an expected signal sequence with the index from 100 to 400 (denoted as “Original signal”, *X_g_*), and CWT with scales *a* as 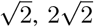 and 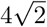 are applied, respectively. Although the details are blurred in the wavelet transformed signals, most peaks and nadirs in CWT(*X_g_*, ·) maintain stability and retain their corresponding locations as in the original signals (e.g., the two green ovals). Furthermore, the shape of the CWT spectrum changes smoothly from a smaller scale value to a larger one, which ensures the success of feature mapping from low-resolution representations to high-resolution ones, and consequently justifies the design of our multi-level algorithm.

**Figure 3:**
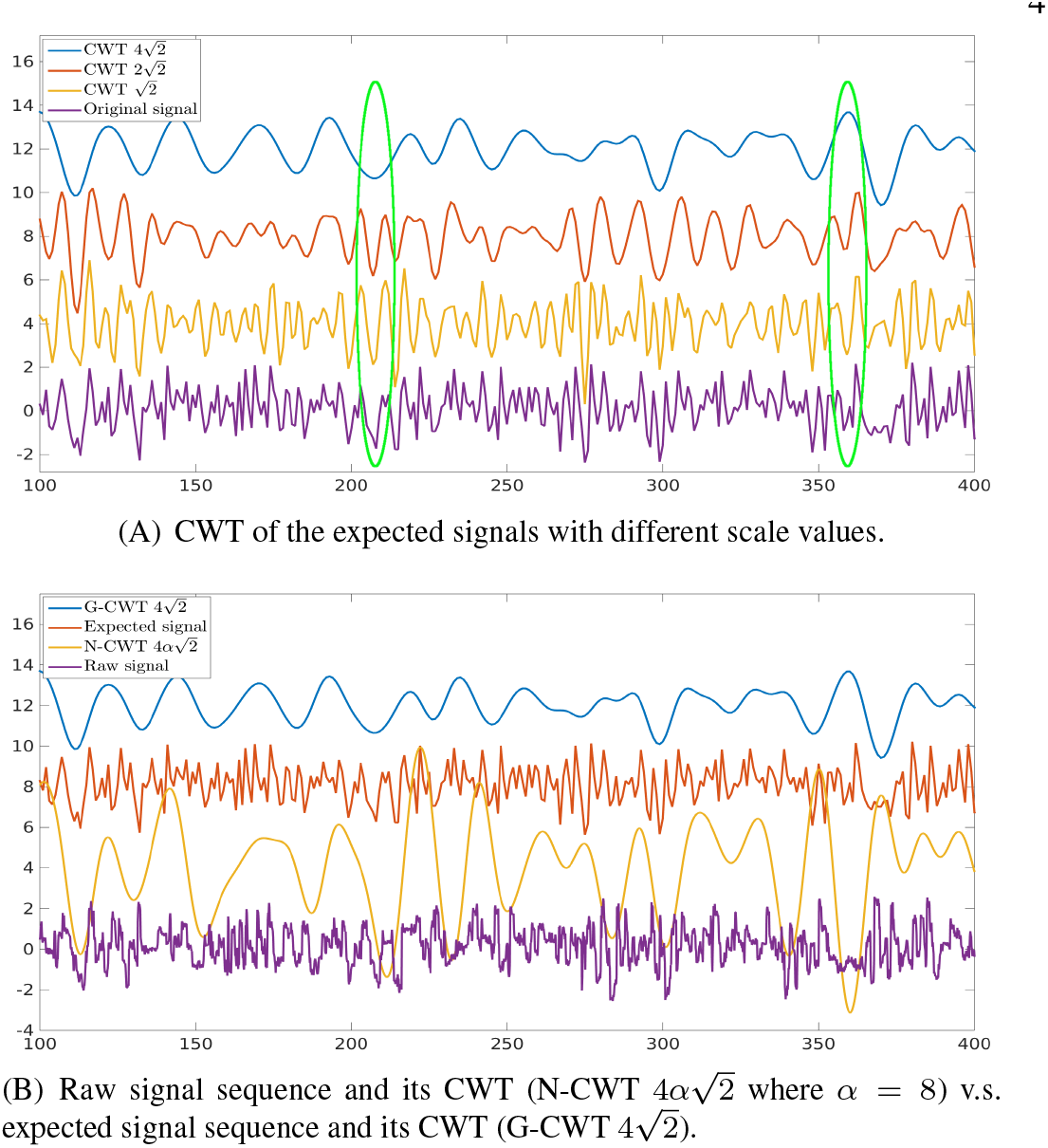
Continuous wavelet transform (CWT) on nanopore signal sequences.

#### 3.1.2 Feature extraction from CWT spectra

The global mapping of two signal sequences with unbalanced lengths due to different sampling rates is a very challenging task, which cannot be handled accurately and rapidly by the existing DTW algorithms. Potentially, re-sampling techniques can be used to alleviate the problem if the degeneration of the accuracy is acceptable. Here we argue that the spectrum analysis based on CWT is a much more natural way to solve the problem.

Fig. 3(B) shows the CWT spectrum comparison of the expected signal sequence (*X_g_*) and the corresponding raw electrical current signal sequence (*X_p_*). Since the lengths of the two signal sequences are one order of magnitude different, where *X_g_* ranges from 100 to 400 and the corresponding *X_p_* ranges from 800 to 3200, we re-scale the index of *X_p_* by 1/8 for the sake of visualization. That is, we apply an additional scale *α* = 8 to the CWT of *X_p_*. It can be seen from Fig. 3(B) that the produced spectrum shape 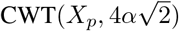 for the raw signal sequence looks quite similar to that of 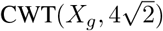 for the expected signal sequence.

We thus propose the following feature extraction procedure to cope with the unbalanced lengths of the raw signal sequence and the expected signal sequence:

1. For signal sequences *X_g_* and *X_p_*, calculate the length ratio *α* = |*X_p_*|/|*X_g_*|, where |*X*| returns the length of *X*;
2. For each scale *a*, obtain the spectra CWT(*X_g_, a*) and CWT(*X_p_, α* · *a*);
3. Normalize CWT(*X_g_, a*) and CWT(*X_p_, α* · *a*) based on Z-score normalization;
4. Extract peaks and nadirs from each spectrum as the feature sequence (we hereinafter call both peaks and nadirs as peaks).

Here, the peaks of the CWT spectrum are extracted as features and will be used for the consequent constrained dynamic time warping (the choice of features will be discussed in Section 4.3.3). Fig. 4 illustrates one round of this procedure. Though the original signal sequences *X_g_* and *X_p_* have significantly different lengths, the numbers of the picked peaks from CWT(*X_g_, a*) and CWT(*X_p_, α* · *a*) are quite similar.

**Figure 4:**
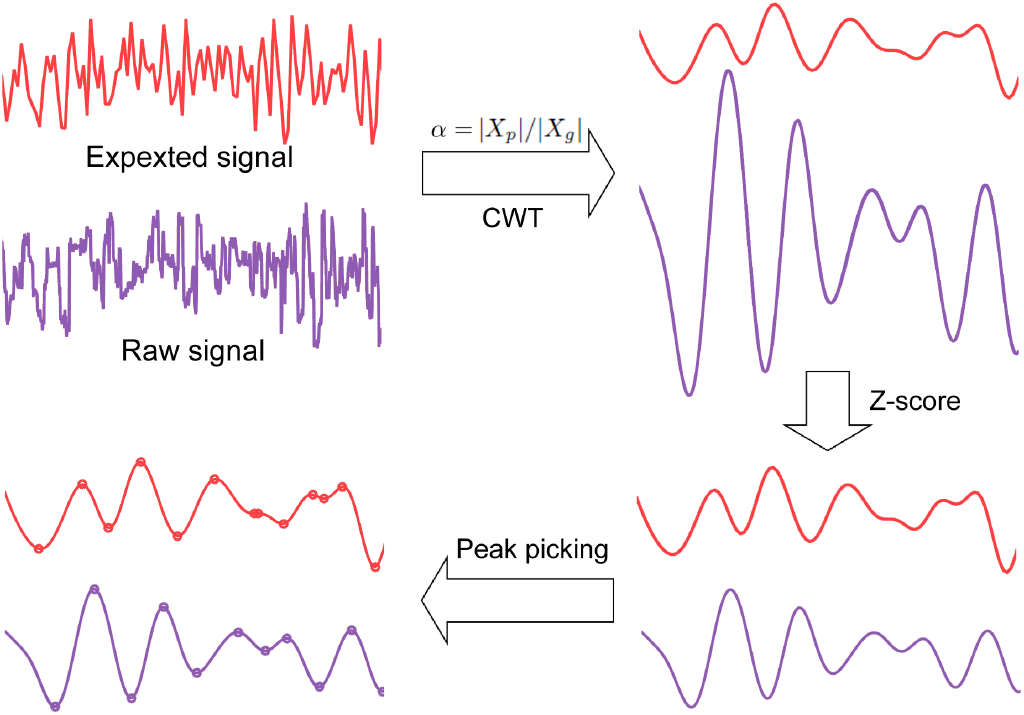
Illustration of the feature extraction procedure from the CWT spectra. The two input signal sequences have unbalanced temporal scales due to different sampling rates. After CWT in consideration of their unbalanced ratio and Z-score normalization, the input signals are converted to CWT spectra with similar shapes. Then the feature sequences are derived by peak picking to make their temporal scales comparable.

### 3.2 Context-dependent Constrained Dynamic Time Warping

The peaks extracted from the CWT spectra are considered as the spectrum features and used in our multi-level dynamic time warping scheme. The main idea is to gradually refine and generate finer warping paths when going from a coarser level to a finer one. We start from the coarsest level *L* where the raw signal sequence is transformed by CWT to 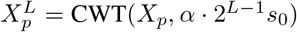 and the expected signal sequence is transformed to 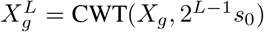. We can then run the original DTW by dynamic programming on the feature sequences (i.e., peaks) from these two transformed sequences. To generate the warping path for the (*L* − 1)th level, we will apply the constrained DTW [23], where the constraints are not predefined, but rather determined by the warping path from the previous level *L*. That is, the warping paths for the (*L* − 1)th level and the *L*th level are not assumed to be the same. In fact, we do not even assume the two paths to have any overlap at all. However, we assume that the warping path for the (*L* − 1)th level is ‘constrained’ by the one for the *L*th level. It should be noted that although the peaks for the (*L* − 1)th level is much more than that for the *L*th level, each peak in the (*L* − 1)th level has a corresponding index interval in the *L*th level. That is, there exists an index *j*, such that this peak in the (*L* − 1)th level resides between the indexes *j* and *j* + 1 at the *L*th level. Our constraint thus requires that each element in the warping path of the (*L* − 1)th level is assumed to be within radius *r* distance from the corresponding element in the path of the *L*th level. Given this context-dependent constraint, the constrained DTW is applied to generate the warping path for the (*L* − 1)th level, which is then used to form the context-dependent constraint for the (*L* − 2)th level. This procedure repeats until the raw signal level is reached, where the final warping path is generated. Section S1 gives the technical details and the pseudo-code for the proposed context-bounded DTW.

Fig. 5 shows one example of the context-dependent constrained DTW with CWT level *L* = 3 and radius *r* = 1. It can be seen that the bounded constraint for each finer level is determined by the warping path from the coarser level, which not only significantly reduces the search space, but also avoids the incorrect mapping at a coarser level being retained at finer levels. This differentiates our algorithm from other approximate DTW algorithms which assume predefined constraints to reduce the search space. Although our algorithm does not require the warping paths at two consecutive levels to overlap, they often do overlap in practice. This is due to the high correlation among the CWT spectra with different scales, which inherits the context of the original signal sequence.

**Figure 5:**
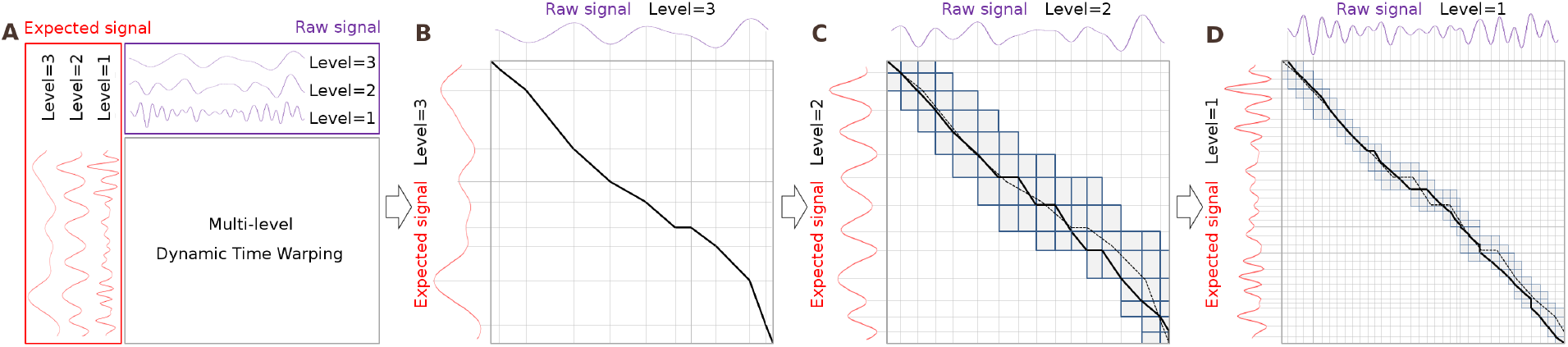
Illustration of multi-level dynamic time warping based on the context-dependent boundary. For illustration purpose, we show here the level up to *L* = 3 and radius parameter *r* = 1. To show the path refinement procedure, dotted lines indicate the warping path of the coarser level, where as solid lines indicate the warping path at the finer level.

### 3.3 Multi-level Continuous Wavelet-based Dynamic Time Warping

In summary, the proposed cwDTW algorithm is shown in Algorithm 1, where *X_p_* is the raw signal sequence and *X_g_* is the expected signal sequence, *L* is the user-defined level, *r* is the searching radius, and *s*_0_ is the CWT base scale; CWT(·) is the continuous wavelet transform defined in Section 3.1.1, PickPeaks(·) is the peak picking procedure described in Section 3.1.2 which returns peak indexes and intensities as two vectors, DTW(·) is the original dynamic time warping algorithm, cDTW(·) is the constrained dynamic time warping [23], and ReMapIndex(·) is the context-dependent constraint generation in Section 3.2.

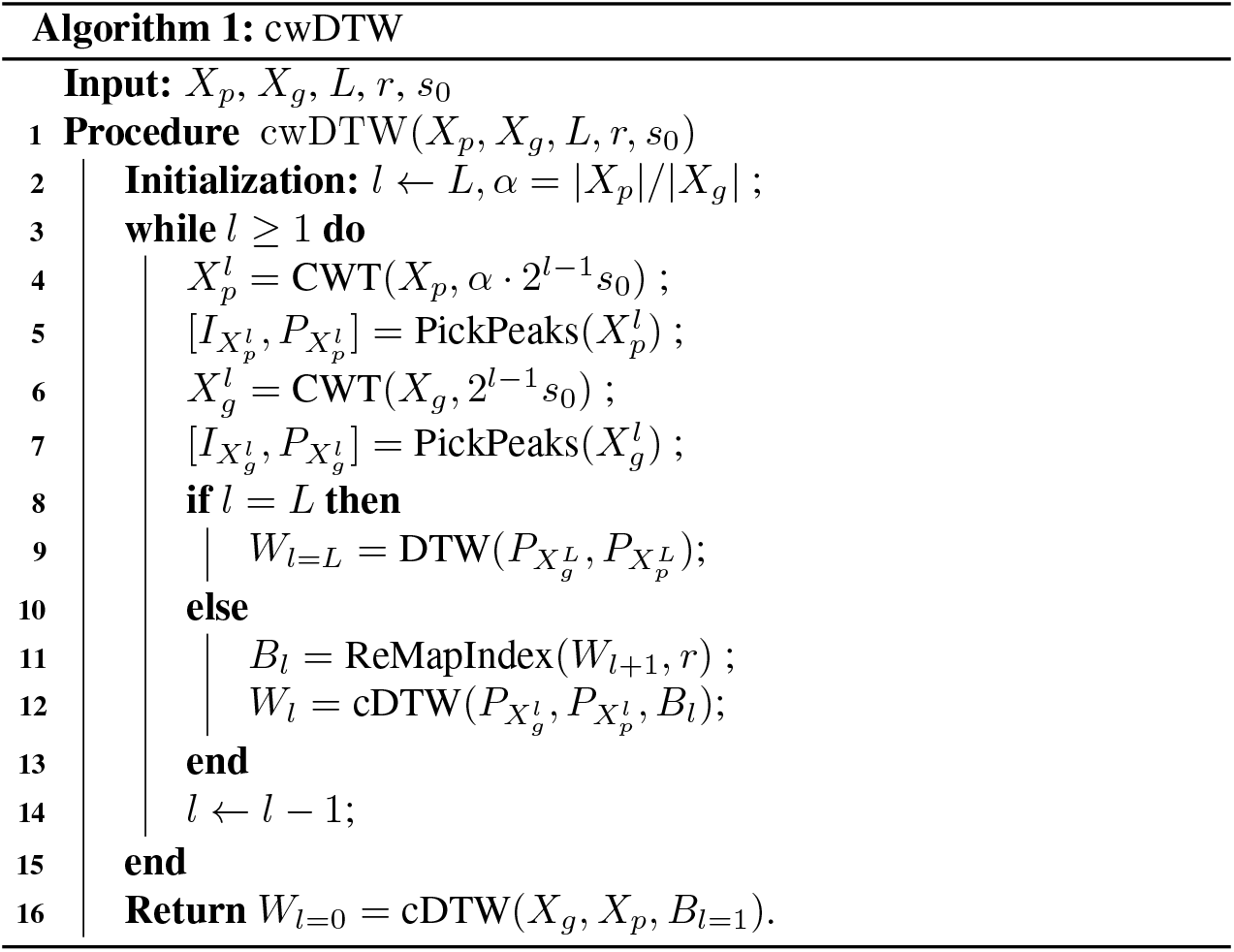

To analyze the runtime and memory complexity of cwDTW, we notice that the operations CWT(·), PickPeaks(·) and ReMapIndex(·) all have *O*(*N*) time- and memory-complexity, where *N* = max {*L*_1_, *L*_2_}. Since the cost matrix of *cDTW* is only filled in the bounded neighborhood of the warping path from the previous level, which grows linearly with a multiplier *r*, cDTW has *O*(*rN*) time- and memory-complexity. The number of picked peaks from the CWT spectrum with the 2^*l*−1^ *s*_0_ scale is upper bounded by 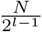. Thus in total, 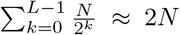 peaks are remapped and considered by constrained DTW. Therefore, the total runtime of Algorithm 1 is 2 · (TIME(cDTW) + TIME(ReMapIndex)) + *L* · TIME(CWT), which gives the time- and memory-complexity of *O*(2*rN* + *LN*), where *r, L* ≪ *N*.

## 4 Experimental Results

We comprehensively evaluated cwDTW on two real nanopore datasets on human and on *Pandoraea pnomenusa*, as well as two benchmark datasets from previous studies [10] which have short sequences with similar lengths. Due to the page limit, we show the results on the two nanopore datasets in this section and refer the readers to Section S2 for the comparison between our algorithm, and FastDTW and PrunedDTW on the two benchmark datasets.

### 4.1 Datasets

Two nanopore sequencing datasets are used in our experiments. The first one is a subset of the public available Human data. This dataset comes from Human chromosome 21 from the Nanopore WGS Consortium [24]. The samples in this dataset were sequenced from the NA12878 human genome reference on the Oxford Nanopore MinION using 1D ligation kits (450 bp/s) with R9.4 flow cells. The nanopore raw signal data in FAST5 format were downloaded from nanopore-wgs-consortium^2^. The reference genome for Human chromosome 21 was downloaded from NCBI^3^. The total number of the generated reads in this dataset is 4530. The average length of the DNA sequences is 7309, whereas the average length of the nanopore raw signal sequences is 68628. The temporal scale ratio between the nanopore raw signal and the corresponding DNA sequence is around 9. We hereinafter denote this dataset as HM4530.

The second dataset is the genome of one bacterial species named *Pandoraea pnomenusa* strain 6399, which was prepared and sequenced by Prof. Lachlan Coin’s lab at University of Queensland. Its reference genome was downloaded from NCBI^4^. The samples were sequenced on the MinION device with 1D protocol on R9.4 flow cells (FLO-MIN106 protocol). The total number of generated reads is 4782. The average length of the DNA sequences is 18590, whereas the average length of the nanopore raw signals is 158772. Thus, the temporal scale ratio between the nanopore raw signal and the corresponding DNA sequence is around 9. We hereinafter denote this dataset as PP4782.

### 4.2 Compared Methods and Evaluation Criteria

Since both datasets have extremely long sequences, and the raw signal sequences and the expected ones have one order of magnitude difference in length, we mainly compared cwDTW with DTW and FastDTW [10]. DTW is the original method that finds the optimal mapping between the two signal sequences by dynamic programming, whereas FastDTW is the state-of-the-art multi-level DTW method which approximates DTW in linear time- and space-complexity. Other representative DTW algorithms, such as PrunedDTW [12], were designed to measure the similarity between two sequences. They implicitly assume that the lengths of the two sequences to be mapped are comparable and thus cannot handle the unbalanced sequences in the two nanopore datasets. Therefore, we only included PrunedDTW in the comparison over the two benchmark datasets (Section S2), which have similar lengths for mapped sequences. Both FastDTW and cwDTW have the radius parameter *r* and the level parameter *L*, and cwDTW has an additional scale parameter *s*_0_ to select the base wavelet scale. We evaluated the performance of cwDTW with different combinations of *s*_0_, *r* and *L* in Section 4.3. All the methods were run on a Fedora25 system with 128Gb memory and two E5-2667v4 (3.2 GHz) cores, each with 8 CPUs.

The accuracy of the warping path *W* generated by an approximate DTW method can be measured by the relative distance difference of *W* with respect to the optimal path *Ŵ* generated by the original DTW [10] (Section 2.2):

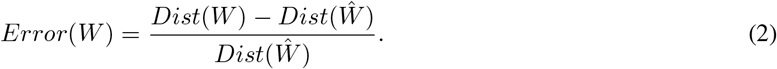

If an algorithm returns a perfect warping path, the error is zero. Note that the error may exceed 100% if the distance of the warping path is more than twice of that of the optimal path. We also calculated the normalized distance of a warping path, *nDist*(*W*), by dividing *Dist*(*W*) by the length of the longer sequence, i.e., the raw signal sequence.

### 4.3 Performance

#### 4.3.1 Performance comparison on nanopore signal mapping with unbalanced lengths

We first compared cwDTW with FastDTW and the original DTW on the HM4530 set. Here both cwDTW and FastDTW have the level set to *L* = 4 and radius set to *r* = 50, and the base wavelet scale for cwDTW is set to 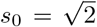. As shown in Fig. 6(A), the distribution of normalized distance of cwDTW is very similar to that of the original DTW. Fig. 6(B) shows the scatter plot between the normalized distance of the original DTW (*x*-axis) and that of cwDTW and FastDTW (*y*-axis). It is clear that the mapping accuracy of cwDTW is very close to the original DTW, whereas FastDTW is far less accurate. In particular, out of the 4530 reads in the HM4530 set, cwDTW produces exactly the same optimal warping path as the original DTW on 3913 reads (86.4%), while on 4393 reads (97.0%) the normalized distance of the path generated by cwDTW is within the 0.005 margin to that of the optimal path. In terms of runtime, it took cwDTW 1406 seconds using a single-CPU (0.31 second on average), whereas FastDTW took 10.7 hours using 16-CPUs in parallel (136 seconds on average) and the original DTW took more than 3 days using 16-CPUs in parallel (916 seconds on average). This implies that cwDTW is about 440 times faster than FastDTW and 3000 times faster than the original DTW.

**Figure 6:**
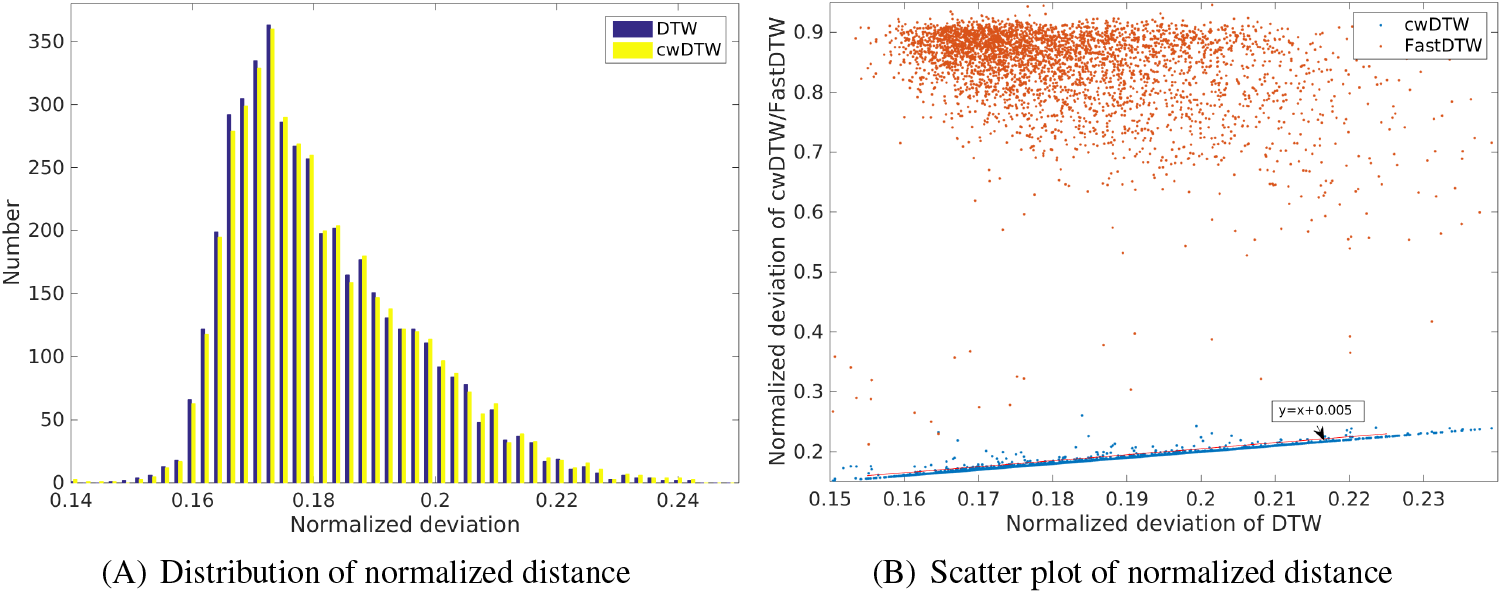
Performance of cwDTW on the HM4530 dataset. (A) Distribution of the normalized distance of cwDTW (in yellow) and the original DTW (in blue). (B) Scatter plot between the normalized distance of the original DTW (*x*-axis) and that of cwDTW (in blue) and FastDTW (in red) (*y*-axis).

We further compared cwDTW with FastDTW under different radius and scale parameter settings on both HM4530 and PP4782 datasets. As shown in Table 1, for different scale parameter values, the mapping error of cwDTW is always lower than 1% if the radius is set to be at least 50. For the PP4782 set, cwDTW can always produce the optimal warping path when the scale is small (e.g., 1 or 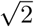) and the radius is large (e.g., 90 and 100). On the contrary, the mapping error of FastDTW remains higher than 200% on both HM4530 and PP4782. Enlarging the radius parameter helps reduce the error for FastDTW, but the effect is marginal.

**Table 1:** Average error of FastDTW and cwDTW with different radius and scale parameter values on the HM4530 and PP4782 datasets. Here we fixed *L* = 4 for both algorithms.

|  | Radius | 10 | 20 | 30 | 40 | 50 | 60 | 70 | 80 | 90 | 100 |
| --- | --- | --- | --- | --- | --- | --- | --- | --- | --- | --- | --- |
| HM4530 | FastDTW | 323% | 311% | 295% | 277% | 261% | 253% | 244% | 228% | 218% | 208% |
|  | cwDTW (2) | 16.8% | 4.8% | 2.1% | 1.1% | 0.6% | 0.4% | 0.3% | 0.2% | 0.2% | 0.2% |
| | cwDTW ($\sqrt{2}$) | 11.5% | 2.6% | 1.1% | 0.6% | 0.4% | 0.3% | 0.2% | 0.2% | 0.2% | 0.1% |
|  | cwDTW (1) | 10.6% | 2.3% | 0.9% | 0.5% | 0.3% | 0.2% | 0.2% | 0.2% | 0.1% | 0.1% |
| PP4782 | FastDTW | 473% | 458% | 431% | 416% | 401% | 395% | 386% | 373% | 365% | 359% |
|  | cwDTW (2) | 30.6% | 9.4% | 3.7% | 1.7% | 0.9% | 0.6% | 0.4% | 0.3% | 0.2% | 0.1% |
| | cwDTW ($\sqrt{2}$) | 22.2% | 4.8% | 1.3% | 0.5% | 0.2% | 0.1% | 0.1% | 0.1% | 0.0% | 0.0% |
|  | cwDTW (1) | 19.1% | 3.8% | 1.0% | 0.4% | 0.2% | 0.1% | 0.1% | 0.0% | 0.0% | 0.0% |

We then conducted a detailed parameter sensitivity analysis of cwDTW to assess the effects of the three parameters, radius, scale and level, on the average error and runtime. In general, cwDTW is quite robust in terms of the average error and runtime with respect to all the three parameters. A high scale parameter (e.g., *s*_0_ = 2) will slightly increase the average error of cwDTW on both datasets (Fig. 7(A) and (E)), while the average error of cwDTW almost remains the same for different level parameters (Fig. 7(B) and (F)). In practice, a radius value of higher than 50 seems to be sufficient. In terms of runtime, different values of the scale parameter do not influence the runtime much (Fig. 7(C) and (G)), whereas a lower level parameter results in higher runtime (Fig. 7(D) and (H)). This is due to the fact that cwDTW runs the original DTW without any constraint for the coarsest level. Thus if L is too small, the number of peaks in the CWT sequences is high, which results in high runtime for the original DTW. Overall, a higher radius value leads to higher runtime. Summing up these observations, a parameter combination of *r* = 60, 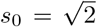 and *L* = 4 gives a practically good tradeoff between error and speed for nanopore signal mapping.

**Figure 7:**
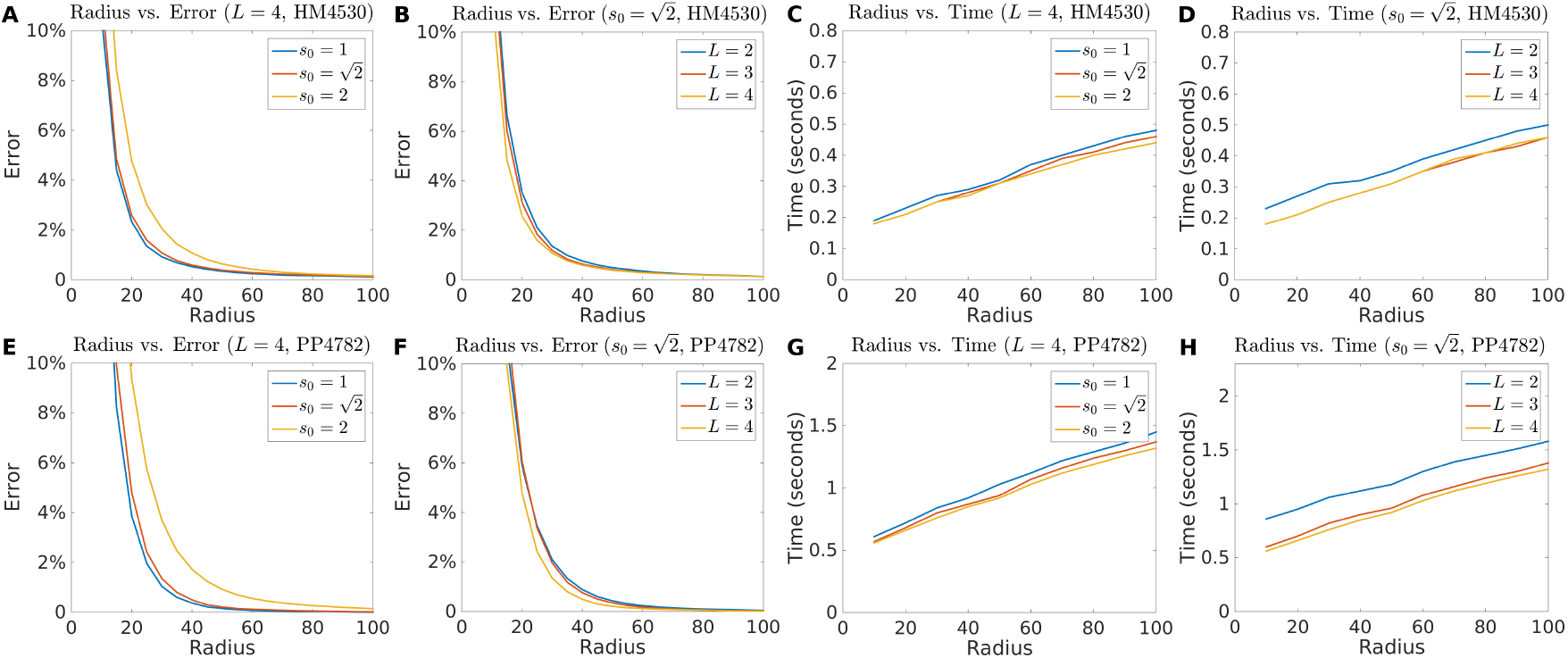
Parameter sensitivity analysis of cwDTW on the HM4530 (upper panel) and PP4782 (lower panel) datasets. (A)&(E): effects of the scale parameter *s*_0_ and radius *r* on the average error over the two sets. (B)&(F): effects of the level parameter *L* and radius *r* on the average error over the two sets. (C)&(G): effects of the scale parameter *s*_0_ and radius *r* on the runtime over the two sets. (D)&(H): effects of the level parameter *L* and radius *r* on the runtime over the two sets.

#### 4.3.2 Performance comparison on signal mapping with similar lengths

One of the main advantages of cwDTW is the ability to handle signal sequences that have orders of magnitude different lengths. This is achieved by the introduction of the scale factor α in consideration of the length ratio during CWT. On the contrary, FastDTW performs coarsening by 2-factor down-sampling, which does not solve the issue caused by length difference and leads to deviations of the warping path. Such deviations will accumulate through iterations and finally corrupt the results.

In order to evaluate the performance of cwDTW on sequences with similar lengths, we created two datasets, HM4530F and PP4782F, from HM4530 and PP4782, respectively. The expected signal sequences in HM4530F and PP4782F are the feature sequences extracted by CWT with scale 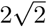 from the expected signal sequences in HM4530 and PP4782, respectively. And the raw signal sequences in HM4530F and PP4782F are the feature sequences extracted by CWT with scale 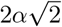 from the raw signal sequences in HM4530 and PP4782, respectively, where α is the temporal ratios in the two sets. Consequently, for HM4530F, the average length of the expected signal sequences is 1558 and that of the raw signal sequences is 1572, whereas for PP4782F, the average length of the expected signal sequences is 3912 and that of the raw sequences is 3957. These two sets thus contain sequences with similar lengths to be mapped.

Table 2 summarizes the average error of FastDTW and cwDTW with different radius and scale parameter values over HM4530F and PP4782F. It is clear that cwDTW still drastically outperforms FastDTW regardless of the parameter settings. When the radius is at least 60, cwDTW can always keep the mapping error to be lower than 1% for both datasets, whereas the error of FastDTW is at least 20%. Further experiments show that to reduce the error of FastDTW to be below 10%, one needs to set the radius parameter to be around 200, which consequently increases the search space and thus runtime dramatically.

**Table 2:** Average error of FastDTW and cwDTW with different radius and scale parameter values on the HM4530F and PP4782F datasets. Here we fixed *L* = 4 for both algorithms.

|  |  | Radius | 10 | 20 | 30 | 40 | 50 | 60 | 70 | 80 | 90 | 100 |
| --- | --- | --- | --- | --- | --- | --- | --- | --- | --- | --- | --- | --- |
| HM4530F | FastDTW |  | 107% | 75% | 59% | 49% | 43% | 38% | 34% | 32% | 29% | 28% |
|  | cwDTW (2) |  | 19.3% | 8.7% | 4.5% | 2.5% | 1.5% | 0.9% | 0.6% | 0.4% | 0.3% | 0.2% |
| | cwDTW ($\sqrt{2}$) | | 17.1% | 6.8% | 2.8% | 1.4% | 0.7% | 0.4% | 0.2% | 0.1% | 0.1% | 0.1% |
|  | cwDTW (1) |  | 15.7% | 5.7% | 2.3% | 0.9% | 0.4% | 0.2% | 0.1% | 0.1% | 0.0% | 0.0% |
| PP4782F | FastDTW |  | 89% | 61% | 47% | 38% | 33% | 28% | 25% | 23% | 21% | 20% |
|  | cwDTW(2) |  | 14.6% | 6.8% | 3.8% | 2.4% | 1.5% | 1.0% | 0.7% | 0.5% | 0.3% | 0.2% |
| | cwDTW($\sqrt{2}$) | | 13.7% | 6.2% | 3.1% | 1.6% | 0.9% | 0.5% | 0.3% | 0.2% | 0.1% | 0.1% |
|  | cwDTW(1) |  | 13.1% | 5.8% | 2.6% | 1.1% | 0.5% | 0.3% | 0.1% | 0.1% | 0.0% | 0.0% |

We further tested cwDTW on two benchmark datasets, Trace and Gunpoint, used in the evaluation of FastDTW [10]. Each of these sets contains sequences with short lengths, ranging from 150 to 275. The comparison results show that cwDTW still significantly outperforms both FastDTW and PrunedDTW in terms of the average error, under all the settings of the radius parameter (Section S2). Taking together all the consistent results from Sections 4.3.1, 4.3.2 and S2, it can be concluded that cwDTW is very robust to the temporal scale, and can handle both balanced and unbalanced cases efficiently and accurately. Two case studies are presented in Section S3 to illustrate how cwDTW can correct errors in the warping path.

#### 4.3.3 Importance of the feature extraction strategy

We further investigated the importance of the feature extraction strategy by peak picking described in Section 3.1.2, for which we compared the proposed strategy with two alternative approaches: equal averaging and peak averaging. Suppose at level *l*, there are *L_l_* peaks. The proposed peak picking strategy uses these *L_l_* peaks as the feature sequence. For equal averaging, the original signal sequence is equally partitioned into *L_l_* bins and the average signal values in these bins are extracted as the feature sequence. For peak averaging, the original signal sequence is partitioned according to the locations of the *L_l_* peaks, and the average signal values in the window composed of the left and right half bins of each peak are used as the feature sequence. Therefore, for all three strategies, the length of the feature sequence is the same.

Figure 8 shows the comparison between FastDTW and cwDTW with three different feature extraction strategies over HM4530F and PP4782F datasets. It is clear that the proposed cwDTW always performs the best regardless of the radius value, followed by cwDTW with the peak averaging strategy for feature extraction. This suggests that the peak signals in CWT contain more stable and useful information than the locally averaged values around the peaks. This is consistent with previous studies which show that CWT can keep a compact and denoised representation of the original signal [18]. The performance of the equal averaging strategy, on the other hand, is sensitive to the choice of the scale parameter and is far less accurate than that of the original cwDTW and the peak averaging strategy. This implies that the peak locations captured by CWT are important to extract useful information. These results justify the use of the peak picking strategy as the feature extraction method in cwDTW.

**Figure 8:**
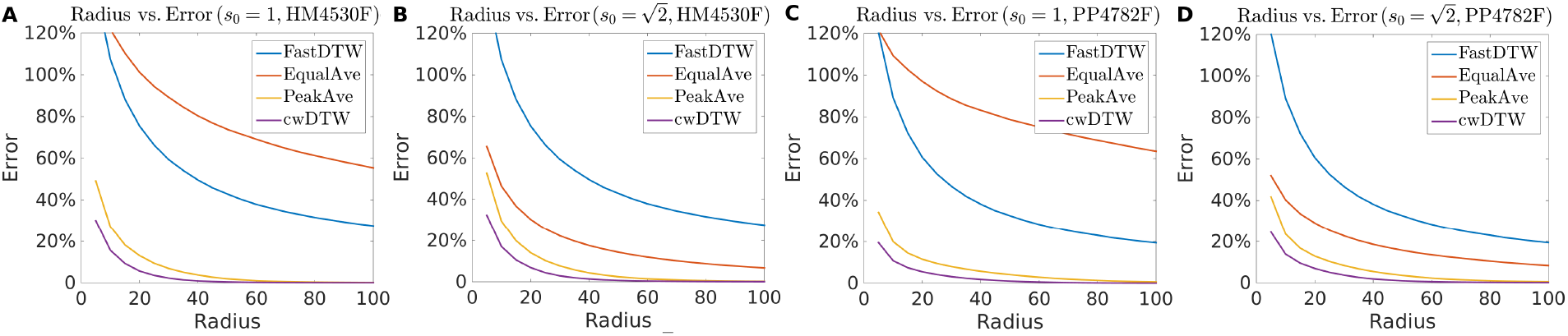
Comparison between FastDTW and cwDTW with three different strategies for feature extraction over the HM4530F and PP4782F datasets. Here we fixed *L* = 4 for all the algorithms.

## 5 Conclusion

We proposed a novel continuous wavelet dynamic time warping algorithm for unbalanced global mapping in nanopore sequencing. The proposed algorithm performs coarsening on the input signal sequences via CWT with different resolutions. Peaks are picked from CWT sequences to form feature sequences. The warping path obtained from a coarser resolution is used to obtain a context-dependent boundary to constrain the warping path at a refiner resolution. Comprehensive experiments on both real nanopore datasets with unbalanced sequences and benchmark datasets with balanced sequences demonstrated the effectiveness and efficiency of cwDTW, which cannot be achieved by the state-of-the-art DTW algorithms. The proposed algorithm provides a powerful tool for various nanopore sequencing tasks, such as base calling, reads mapping, variant identification, and methylation detection. In addition, the generic nature of cwDTW makes it a useful method for mapping temporal sequences of biological, video, audio and graphical data.

## Acknowledgement

We thank Minh Duc Cao, Lachlan J.M. Coin, Louise Roddam, and Tania Duarte for providing the nanopore sequencing data for the *Pandoraea pnomenusa* sample.

## Appendix

### S1 Context-dependent Bounded Dynamic Time Warping

It should be noted that all the peaks are directly extracted from the CWT of the signals, which is the main difference between our algorithm and other DTW variants [10, 16, 17]. Different scales of CWT will result in peaks on different levels. In our algorithm, given the base scale *s*_0_ and the level *L*, for *l* ⩽ *L*, the lth level scale is defined as *s_l_* = 2^*l*−1^ *s*_0_. Consequently, the *l*th CWT spectra for the raw signal sequence and the expected signal sequence are defined as 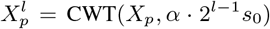 and 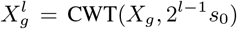, respectively. The peak extraction defined in Section 3.1.2 is denoted as operation [*I_X^l^_, P_X^l^_*] = PickPeaks(*X^l^*), where *P_X^l^_* is the vector of peak intensities and *I_X^l^_* is the vector of peak indexes in the sequence *X^l^*.

Our algorithm starts with the coarsest level *L*. For 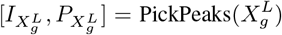 and 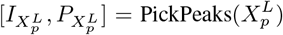, there is no boundary that can be used for mapping and the feature sequences are short. Thus, the original DTW is applied to map 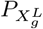 and 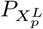 (as illustrated in Fig. 5(B)). Here, we denote 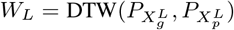 as the path generated by DTW from the *L*th level feature sequences. For each *w_L_* = (*i, j*) element in *W_L_*, a mapping related to *X_g_* and *X_p_* can be obtained by remapping 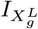 and 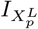. For the (*L* − 1)th level, we constraint the search space according to the warping path from the *L*th level. In general, for any arbitrary level *l*, the feature sequences 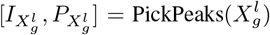 and 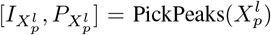 can be mapped by DTW constrained by the warping path, *W*_*l*+1_, from the (*l* + 1)th level. The constraint is determined as follows:

1. Given *W*_*l*+1_, remap each *w*_*l*+1_ = (*i, j*) to the original resolution sequences *X_g_* and *X_p_* to gain the mapping 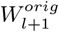;
2. For each 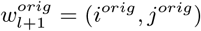, search in 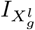 to find index *i*′ such that 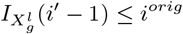 and 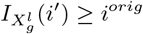; and find index *j*′ such that 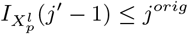 and 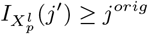;
3. Form a tentative path, 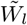, for level *l* by setting 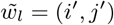;
4. Interpolate between (*i*′ − 1, *j*′ − 1) and (*i*′, *j*′), i.e. fill the index gap in 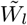 to create an interpolated warping path 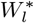;
5. Generate the search boundary from 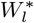 by extending each element (*i, j*) in four directions with radius *r* to create an area *i* − *r*, *i* + *r, j* − *r, j* + *r* (as illustrated in Fig. 5(C) and (D)).

That is, the warping path at level *l* is searched within a narrow boundary along the warping path at level *l* + 1, which makes constrained DTW very efficient. On the other hand, we do not assume the warping paths at levels *l* and *l* + 1 to have any overlap, which prevents the mapping error to be propagated.

### S2 Performance Comparison on Benchmark Datasets with Similar Lengths

Two datasets from the UCRArchive^5^ are used to compare the performance of cwDTW, PrunedDTW^6^ and FastDTW. The two datasets are the Trace dataset and the Gunpoint dataset, which were also used to evaluate FastDTW [10]:

**Figure S1:**
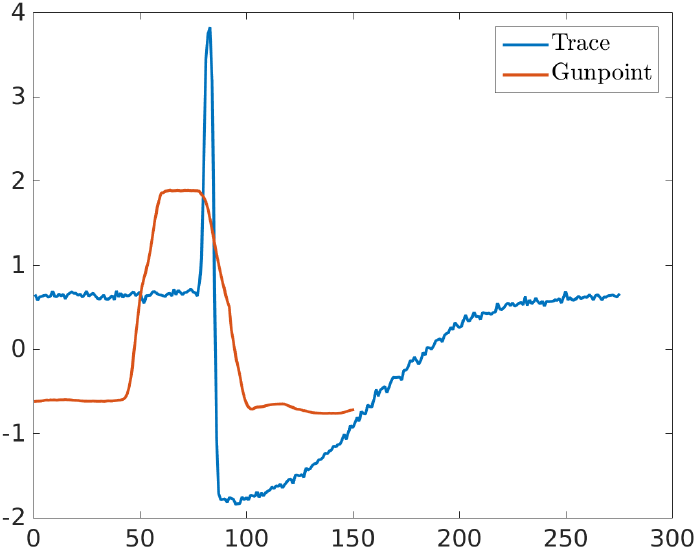
Illustration of the representative time series sequences in the Trace (in blue) and the Gunpoint (in red) datasets.

**Trace** contains 200 time series sequences. This dataset contains real instrumentation data from a nuclear power plant with several types of simulated transient events inserted that simulate instrumentation failure. Each time series sequence has a length of 275 points. Time series in this dataset have similar overall shapes, as illustrated as the blue curve in Fig. S1.

**Gunpoint** contains 200 time series sequences. Each time series sequence has a length of 150 points. The dataset contains two classes, a gun being drawn from a holster and a gun being pointed. The movements are represented by a time series of the hand’s x-axis position over time. An illustration of a representative sequence is given as the red curve in Fig. S1.

In this experiment, we randomly selected one of the sequences in each dataset as the reference sequence and used the others as the query sequences. According to the shapes of the sequences of the two datasets, we roughly set the scale parameter *s*_0_ of cwDTW to 100 for the Trace dataset and 60 for the Gunpoint dataset. Since the sequences in these datasets are very short, we used *L* = 1.

**Figure S2:**
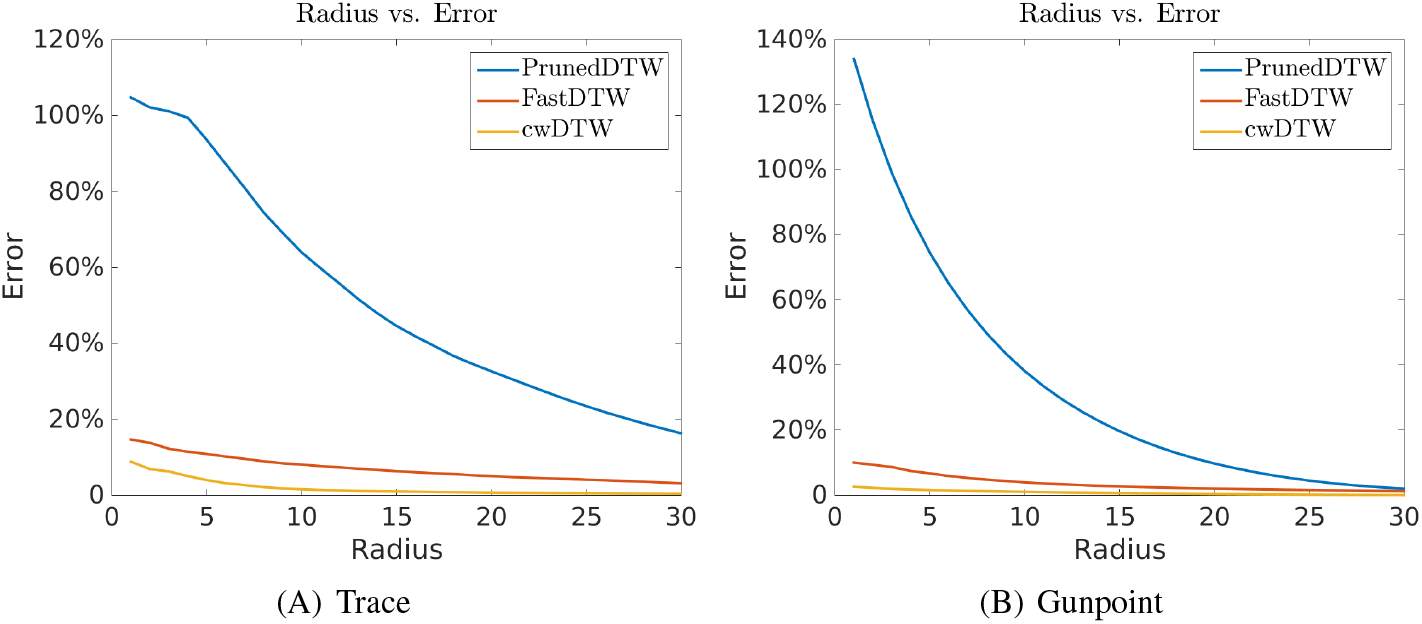
Performance comparison in terms of the relative distance error of cwDTW, FastDTW and PrunedDTW on the Trace and Gunpoint datasets. The *x*-axis is the radius parameter and the *y*-axis is the relative distance error to the original DTW.

Fig. S2 shows the comparison results of cwDTW, PrunedDTW and FastDTW with different settings of the radius parameter. It is clear that cwDTW is always the best method on both datasets in terms of the relative distance error, regardless of the radius parameter. FastDTW performs significantly better than PrunedDTW. In particular, when the radius is at least 15, cwDTW can reach an error rate that is very close to zero. Therefore, cwDTW performs consistently well on mapping short sequences with similar lengths.

### S3 Case Studies

Here we show two case studies to investigate the reason for cwDTW’s superior performance. The first case study is a test case from the Trace dataset used by FastDTW [10] and the second one is a test case from the HM4530^7^ dataset. Here, we fixed *L* = 4, and *r* = 10 for the first case and *r* = 50 for the second one, for both FastDTW and cwDTW 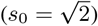.

**Figure S3:**
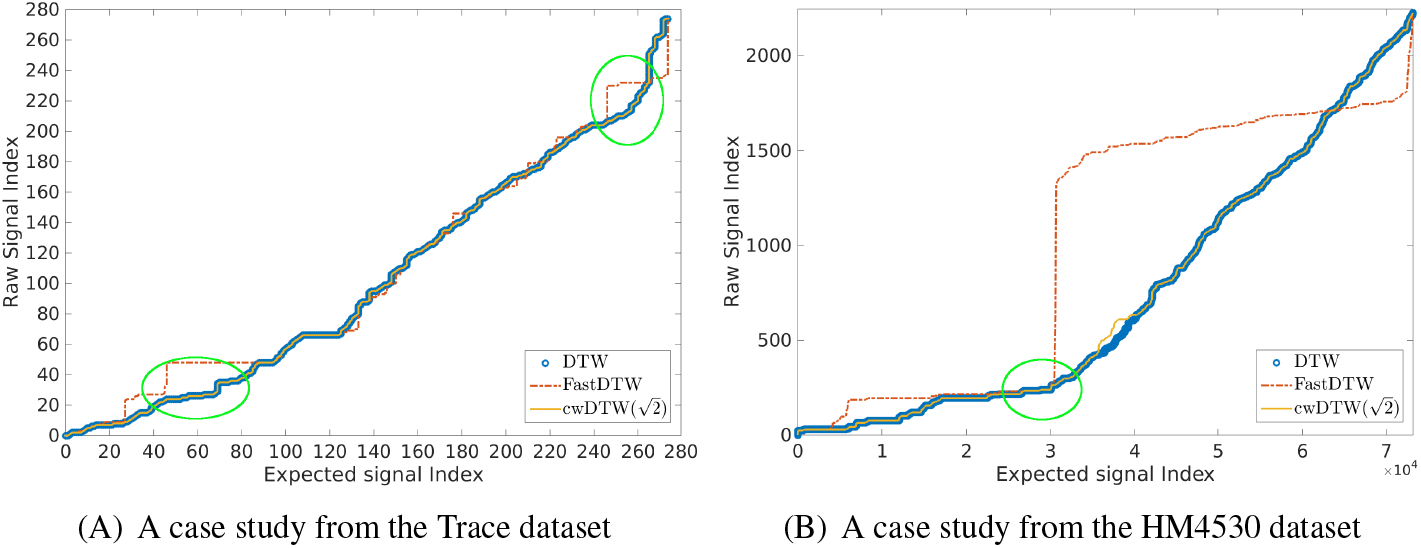
Two case studies. The *x*-axis is the index of the expected signal sequence and the *y*-axis is the index of the raw signal sequence. The blue, red and orange curves represent the warping paths returned by the original DTW, FastDTW and cwDTW, respectively.

From Fig. S3, we can find that the warping path generated by cwDTW is very close to that by the original DTW. As shown in Fig. S3(A), for the first case, the warping path generated by cwDTW is exactly the same as the one produced by the original DTW, whereas the warping path generated by FastDTW has two regions with clear deviations (marked in green ovals). For the much longer sequences shown in Fig. S3(B), the warping path generated by cwDTW still overlaps with the one by DTW in most of the cases, whereas the one generated by FastDTW differs significantly from the optimal path. From the location marked by the green oval in Fig. S3(B), the warping path of FastDTW starts deviating from the optimal path and this deviation propagates. This is due to the fact that FastDTW uses a down sampling strategy and thus the points with important information might be lost. On the other hand, since the peaks of the CWT spectra are context-dependent, it is possible for cwDTW to correct the mapping errors as the resolution becomes finer. This is demonstrated by the deviation in the warping path generated by cwDTW at index around 40000 in the *x*-axis in Fig. S3(B), where cwDTW corrects the mapping error so that the following path converges back to the optimal one.

## Footnotes

1 An expected signal sequence is obtained by applying sliding windows of *k*-mers (here *k* = 5) on the DNA sequence and using the expected electrical current value for each *k*-mer according to the pore model.

2 http://s3.amazonaws.com/nanopore-human-wgs/rel3-fast5-chr21.part03.tar

3 https://www.ncbi.nlm.nih.gov/nuccore/NC_000021

4 https://www.ncbi.nlm.nih.gov/nuccore/JTCR01000000

5 http://www.cs.ucr.edu/~eamonn/time_series_data/

6 PrunedDTW is a recent DTW algorithm and the code can be found at http://sites.labic.icmc.usp.br/prunedDTW/.

7 Data ID: c1bbcb7c-dc57-4469-9a5d-2cc486d5edd5.

